# The structurome of a *Clostridium difficile* phage and the remarkable accurate prediction of its novel phage receptor-binding protein

**DOI:** 10.1101/2021.07.05.451159

**Authors:** Ahmed S. A. Dowah, Guoqing Xia, Ali Abdul Kareem Ali, Anisha M. Thanki, Jinyu Shan, Andy Millard, Bent Petersen, Thomas Sicheritz-Pontén, Russell Wallis, Martha R. J. Clokie

## Abstract

As natural bacterial predators, bacteriophages have the potential to be developed to tackle antimicrobial resistance, but our exploitation of them is limited by understanding their vast uncharacterised genetic diversity^1,2^. Fascinatingly, this genetic diversity reflects many ways that phages can make proteins, performing similar functions that together form the familiar phage particle. Critical to infection are phage receptor-binding proteins (RBPs) that bind bacterial ‘receptors’ and initiate bacterial entry^3^. Here we identified and characterised Gp22, a novel RBP for phage CDHS-1 that infects pathogenic *C. difficile*, but that had no recognisable RBPs. We showed that Gp22 antibodies neutralised CDHS-1 infection and used immunogold-labelling and transmission electron microscopy to identify their location on the capsid. The Gp22 three-dimensional structure was resolved by X-ray crystallography revealing a new RBP class with an N-terminal L-shaped α-helical superhelix domain and a C-terminal Mg^2+^-binding domain. The findings provide novel insights into *C. difficile* phage biology and phage-host interactions. This will facilitate optimal phage development and future engineering strategies^4,5^. Furthermore, the AlphaFold2-predicted Gp22 structure, which was strikingly accurate, paves the way for a structurome based transformation and guidance of future phage studies where many proteins lack sequence homology but have recognisable protein structures.

## Introduction

Bacteriophages, or phages, are viruses that specifically infect and kill bacteria which makes them suitable for development into highly precise antimicrobials. They are the most abundant life form known, but because of their vast genetically diversity, only a small proportion of potentially useful therapeutic phages have genes with predicted functions. This has significantly limited our understanding and exploitation of them as novel antimicrobials. The newly developed machine learning method AlphaFold2 is likely to influence many spheres^6,7^. Because phages have so much novelty^8^ the influence on this field is particularly significant to identify proteins that have structural non-sequence-based relationships.

One therapeutically interesting phage is CDHS-1 that kills the hypervirulent *C. difficile* ribotype 027. *Clostridium difficile* is problematic as it causes severe gastrointestinal disease in humans^9^. *C. difficile* infection (CDI) is often associated with antibiotic treatment, disruption of gut microbiota and overgrowth of *C. difficile^10,11^*. Treatment is difficult as *C. difficile* is naturally resistant to most antibiotics, so alternative treatments for CDI are urgently required^12^. Phages are attractive for this as they are generally species-specific causing minimal disruption to gut microbiota and they are able to self-amplify at infection sites^13^.

CDI phage therapy with CDHS-1 has been used successfully in animal models to reduce bacterial carriage and increase survival following infection. Although we have obtained a lot of phenotypic data on the efficacy of this phage to kill *C. difficile*, little mechanistic information is known on how the phage interacts with the bacterial host. Indeed, standard homology searches only identify a putative function for 15 of the 53 protein encoding genes. To overcome this, we predicted all protein structures by employing the folding algorithm AlphaFold2.

A key aspect of developing phage therapeutics is to have a detailed understanding of the physical interactions between phage Receptor Binding Proteins (RBPs) and their bacterial host receptors that initiate successful infection^14^. RBPs are vital determinants of the phage host range^15^ and are located at the end of the phage tail, in order to bind to ligands (receptors) on the bacterial surface^16^. Phage RBPs are structurally diverse, and many bacterial cells surface molecules can act as receptors including surface proteins, carbohydrates, or wall teichoic acids for Gram positive bacteria^2^. This information is only known for a few phages, which limits our ability to construct phage cocktails and engineer specific target ways^17,18,3^. Currently, the only robust way to identify phage receptor binding proteins (RBPs) is through biochemical and structural approaches. Recent advances in machine learning algorithms for protein structure prediction could help inform such studies and revolutionise the time to obtain crystal structures if the predicted model is identical enough to the experimental one to allow phasing of the crystal diffraction pattern through molecular replacement.

Although sequence similarity is often limited between phages, comparative genomic analysis of phages infecting Gram-positive bacteria including *B. subtilis, L. lactis, L. monocytogenes* and *S. aureus*, has shown that genes encoding phage tail proteins are often located in a tail module between *tmp* gene encoding the tape measure protein (TMP), and the holin and endolysin proteins^19^. Hence, we focused our studies in this area of the genome.

Relatively few Firmicutes phage RBPs have been characterized, and the best known examples are lactococcal phages p2, bIL170, and TP90^20,21,22^ and *Staphylococcus aureus* phage φ11^23^. Despite several *C. difficile* phages being isolated^24,12^ their RBPs have not been identified and little is known about their bacterial receptors. The dominant S-Layer proteins (mainly constructed of S-layer protein A SlpA) of *C. difficile* are likely phage receptors. Recently, Kirk et al. showed that the S-layer protein SlpA acts as a receptor for a product known as Avidocin-CDs, a phage RBP fused with a R-type bacteriocin scaffold to specifically kill *C. difficile* ribotype 027 strains^25^. This RBP has no sequence similarity to any genes encoded within CDHS-1 and has no published data on the structure.

Here we report the identification of the RBP from CDHS-1 and describe its three-dimensional structure as determined by X-ray crystallography and very accurately predicted by AlphaFold2. Both the predicted structure and the crystallography data show that Gp22 (ALY06965.1) is a new class of RBP; it is a stable homodimer consisting of an N-terminal L-shaped α-helical superhelix domain and a C-terminal β-sandwich domain. Reciprocal contacts between the monomers are mediated via an α-helical hairpin that projects from the long arm of each super helical domain to form a four-helix bundle. Interestingly, protomers of the dimer are asymmetrical with the short arm of one polypeptide interacting with its partner.

This is the first study to use AlphaFold2 to generate the predicted structure for a novel phage protein. The striking accuracy (Figure 4e) suggests that current advances in machine learning based protein structure predictions may revolutionise the speed by which such novel structures can be identified and confirmed using structural biology approaches. This work will form the basis of a mechanistic understanding of host recognition for these phages and identifies a novel class of RBPs thus expanding our comprehension of ‘Phage receptor binding protein space’ which we anticipate will be further expanded by the application of AlphaFold2 directed structural studies. These findings further encouraged us to predict all structures for the whole phage proteome, which we term as the complete predicted ‘structurome’.

## Results

### Characterisation of the tail module

To identify the receptor binding proteins (RBPs), the putative tail module was analysed and four likely tail fiber open reading frames; Gp18 (ALY06961.1), Gp19 (ALY06962.1), Gp21 (ALY06964.1), and Gp22 (ALY06965.1) were identified. The *in-silico* sequence prediction tools HHpred and Phyre2^26,27^ revealed that Gp18 is homologous to *B. subtilis* phage SPP1 Gp19.1, a distal tail protein (Dit) (PDB 2X 8K, FigureS1), that forms the central hub of the SPP1 baseplate, thus Gp18 is most likely the CDHS-1 Dit.

HHpred analysis showed that CDHS-1 Gp19 (ALY06961.1) is homologous to *L. Monocytogenes* phage A118 Gp18 (NP_465809.1) from (PDB 3GS9) (FigureS1) and Mu Gp44 (P08558), both tail-associated lysine-like proteins (Tal). The function of these proteins is to help phages inject genetic material into the bacteria by degrading peptidoglycan^19^. Consistent with this, a Gp19 (ALY06961.1) blastp analysis identified an endopeptidase domain (residues 7-378).

CDHS-1 Gp21 (ALY06964.1) is homologous to the ORF48 (NC_002747.1) upper baseplate protein N-terminal domain (BppU) (PDB 3UH8) of phage TP901-1 of *L. lactis* (FigureS1). This protein links the RBP to the central baseplate core so it likely has a similar role in CDHS-1.

Finally, *in silico* Gp22 (ALY06965.1) analysis showed no similarity to any protein structures in the Protein Data Bank (PDB)^28,29^, suggesting it could be a novel RBP. Of note, Gp22 (ALY06965.1) is homologues to *C. difficile* phages phiCD38.2 and phiCD146 which only differ by a single amino acid, and to phiCD111 (82% similarity). There are no three-dimensional structures resolved for any of these proteins. Interestingly, the C-terminal 89 residues are less well-conserved (64% match) than the N-terminal 529 residues (86% match) (FigureS2). This is consistent with well-studied RBPs in Lactococcal phages where receptor-binding sites are within the C-terminal (non-conserved) RBP region.

### Visualising the tail module and the complete structurome with AlphaFold2 Prediction

Although we were excited by the newly determined crystal structure, it took several years to obtain, challenging the applicability of this approach to determining the structures present within the vast phage protein landscape. Although there is extremely high sequence diversity - phages look the same and thus are constrained by functionality^30^. This led us to investigate whether the protein structure prediction algorithm AlphaFold2 could accurately predict novel phage structures, despite their lack of homology to proteins of known structures. To test this, we used AlphaFold2 to predict the structure of all 53 CDHS-1 proteins^6^ (Figure1). The predicted structures are ranked by the mean plDDT (predicted Local Distance Difference Test) confidence score, with an average of 84% (Figure1). Remarkably 90% of the proteins had good confidence scores (Supplementary Table 2), with 40% having very high confidence scores and only four proteins have low scores and one conserved hypothetical protein has an unusable prediction.

**Figure 1:**
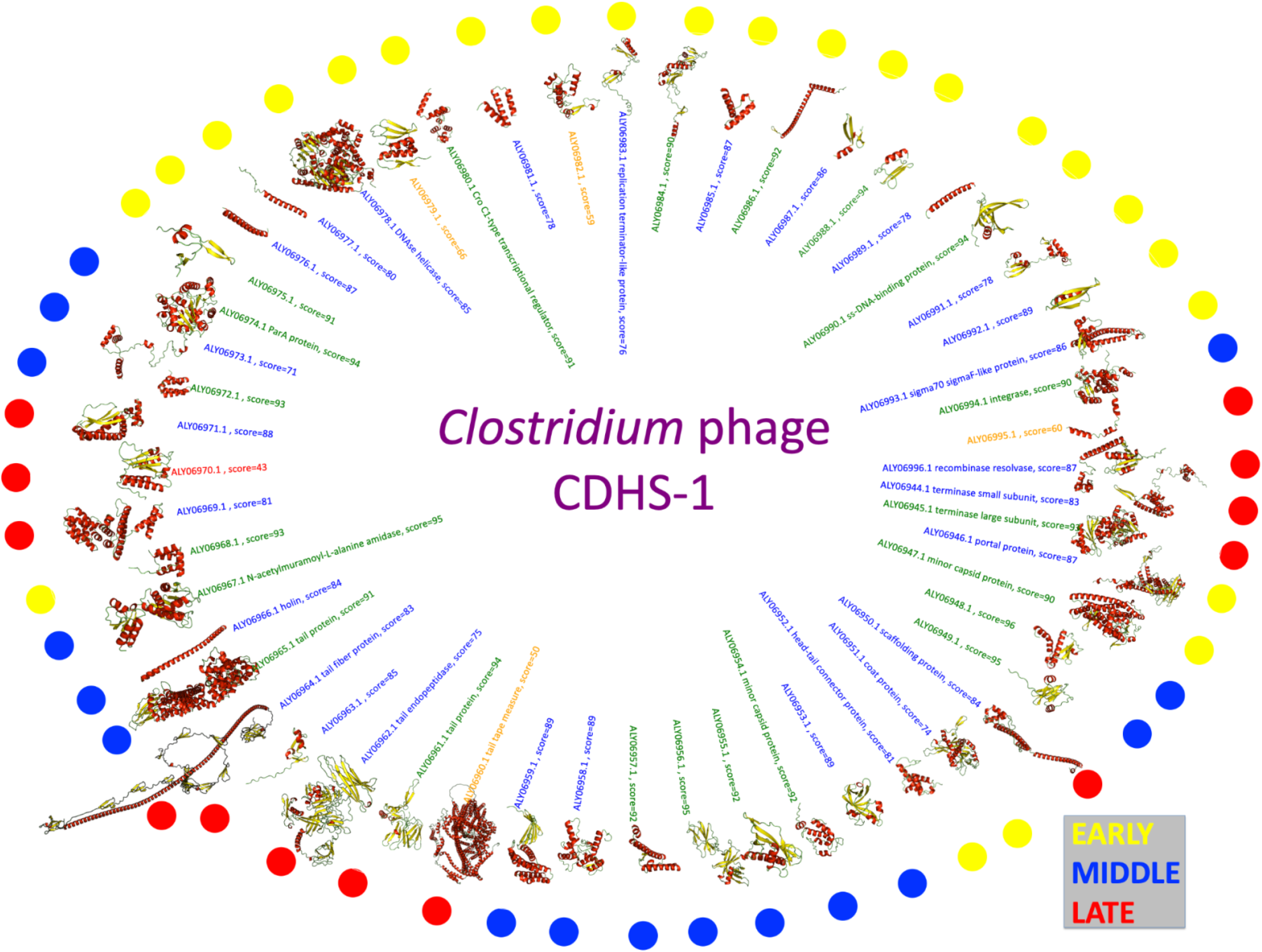
AlphaFold2 predictions of all structures of the whole phage proteome ‘structurome’: All the structures of the 53 CDHS-1 proteins predicted using AlphaFold2 with average confidence scores 84%; this represented by ~59.9% of the proteins have good confidence scores, and 39.6% very high confidence scores. Only 4 proteins were seen to have low scores and one conserved hypothetical protein has an unusable prediction. The circular markers are depicting the temporal expression phase, where yellow marked genes are expressed early, blue middle and red late.

To determine if there is a correlation between the predicted structures and the temporal expression of the genes which encode them^13^, the predicted proteins were divided into three categories - early, middle and late - based on their expression profiles calculated from our previous work (Figure1). Genes expressed at the early phase (yellow) encode proteins that are relatively small and simple in structure, whereas the proteins encoded by late genes (red) are larger and more complex. This is likely to be attributed to the phage take-over strategy where early genes encode for proteins that interfere with bacterial metabolism as they repurpose it for their own purpose^31^, such proteins will interact with polymerases and therefore don’t have complex structures^32^. In contrast, late genes that encode the proteins that make up the phage capsid are more complex. Most of the early expressed proteins are relatively small, apart from the predicted portal, helicase (ALY06978.1) and minor capsid protein. In the early gene stretch three hypothetical proteins have simple linear shapes, but no sequence similarity (ALY06977.1, ALY06976.1 and ALY06989.1), they may either be forming complexes within themselves or with bacterial encoded proteins.

For the proteins encoded by late genes, of particular interest, are the proteins involved in phage structure such as the tape measure proteins (TMP) that have a predicted structure that is clearly similar to known TMPs that were defined using genetic studies^33^. However, no TMPs have been crystalized yet. This illustrates the power of AlphaFold2 to revolutionise our interpretation of phage genomes by providing structure-based predictions.

More importantly, AlphaFold2, accurately predicted the structure of Gp18 (ALY06961.1) and Gp19 (ALY06962.1) and for Gp21(ALY06964.1) only the N-terminal of Gp21(ALY06964.1) is similar when the protein predicted using HHpred and Phyre2 (FigureS1). And above all, predicted Gp22 (ALY06965.1) with highly striking accuracy. As an interesting observation we can see that all proteins encoded by late genes have a complex structure such as Gp18 (ALY06961.1), Gp19 (ALY06962.1), Gp21(ALY06964.1) and Gp22 (ALY06965.1). The reason behind the complexity of these proteins might be that these proteins are involved in highly complicated processes such as bacterial phage interactions, which requires for example more than one copy of the RPBs to interact with the bacteria, depending on the type of receptors that phage is attached to. More copies of RBPs when the receptors on the bacteria are carbohydrates^1^.

### Phage neutralization using anti-Gp22 polyclonal antibodies

To determine if Gp22 (ALY06965.1) is essential for phage infection, polyclonal antibodies were raised against recombinant Gp22 (ALY06965.1) and used in neutralization assays of CDHS-1 to C. *difficile* strain CD105LC1. Antibodies were also raised against the putative baseplate protein Gp21 (ALY06964.1) as a control. In infection assays, pre-incubation with anti-Gp22 (ALY06965.1) serum led to complete inhibition of phage infection, up to a dilution of 1:10000 (Figure 2b) demonstrating it is essential for successful infection of *C. difficile* by CDHS-1. In contrast, pre-incubation of anti-Gp21 (ALY06964.1) serum had no impact on CDHS-1 infection (Figure 2a).

**Figure 2:**
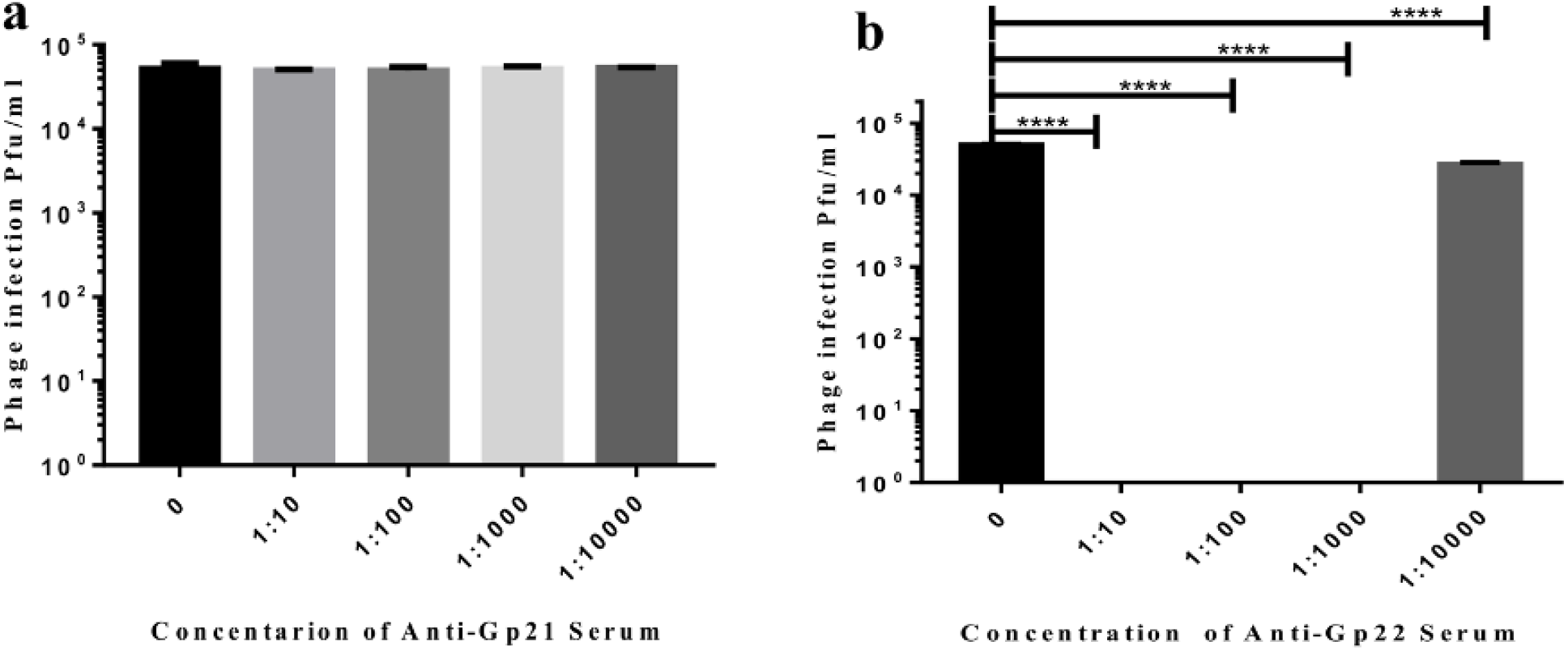
Characterisation of Gp21 and Gp22 (ALY06965.1) proteins in CDHS-1: CDHS-1 was pre-incubated with polyclonal antibodies against the tail proteins Gp21(ALY06964.1) and Gp22 (ALY06965.1). The phage was tested for its ability to infect CD105LC1 using spot tests with serial 10-fold dilutions. (A) And (B) show the plaque-forming units (Pfu/ml) of CDHS-1 on CD105LC1 after incubation with different concentrations of polyclonal antibodies against Gp21(ALY06964.1) and Gp22 respectively. The result in (A) shows no significant differences between the positive control (gray column) and the negative control (black column) when anti-Gp21(ALY06964.1) was used. (B) Represents the inhibition of phage CDHS-1 infection on CD105LC1 strain, using different concentrations of anti-Gp22 (ALY06965.1) serum. The assay was conducted with three biological repeats each with three technical repeats. Error bars represent means ± standard deviations (SD, n = 3). Statistical differences were calculated by two-way ANOVA.

### Recombinant Gp22 binds to *C. difficile* CD105LC1

To confirm that Gp22 (ALY06965.1) is the RBP, recombinant Gp22 (ALY06965.1) was produced as a GST fusion by expression in *E. coli* and *C. difficile* CD105LC1 cells were incubated with glutathione-Sepharose 4B beads coated with GST alone or GST-tagged recombinant Gp22 (ALY06965.1). Confocal microscopy under phase contrast revealed that only bacterial cells attached to beads coated with GST-tagged Gp22 (ALY06965.1) (FigureS3a) were visible, not those coated with GST alone (FigureS3b), indicating that Gp22 (ALY06965.1) binds directly to *C. difficile* cells. Increasing the concentrations of recombinant Gp22 incubated with *C. difficile* before adding CDHS-1 inhibited phage adsorption relative to dose, further confirming that this protein is the RBP (FigureS3c).

### TEM Localisation of Gp22 and Gp21

To identify the precise locations of Gp22(ALY06965.1) and Gp21(ALY06964.1) on the phage particle, CDHS-1 pre-incubated with either rabbit anti-Gp21 or anti-Gp22 (ALY06965.1) sera was incubated with goat anti-rabbit secondary antibodies coupled to gold colloids. The gold-labelled antibodies were seen as black spots under TEM, showing the location of each protein, Gp21(ALY06964.1) and Gp22 (ALY06965.1) are located on the tail baseplate (Figure 3a and b). As expected, only a random scattering of black spots was observed on the grids containing CDHS-1 incubated with the pre-immune sera as a negative control (Figure 3d and 3e).

**Figure 3:**
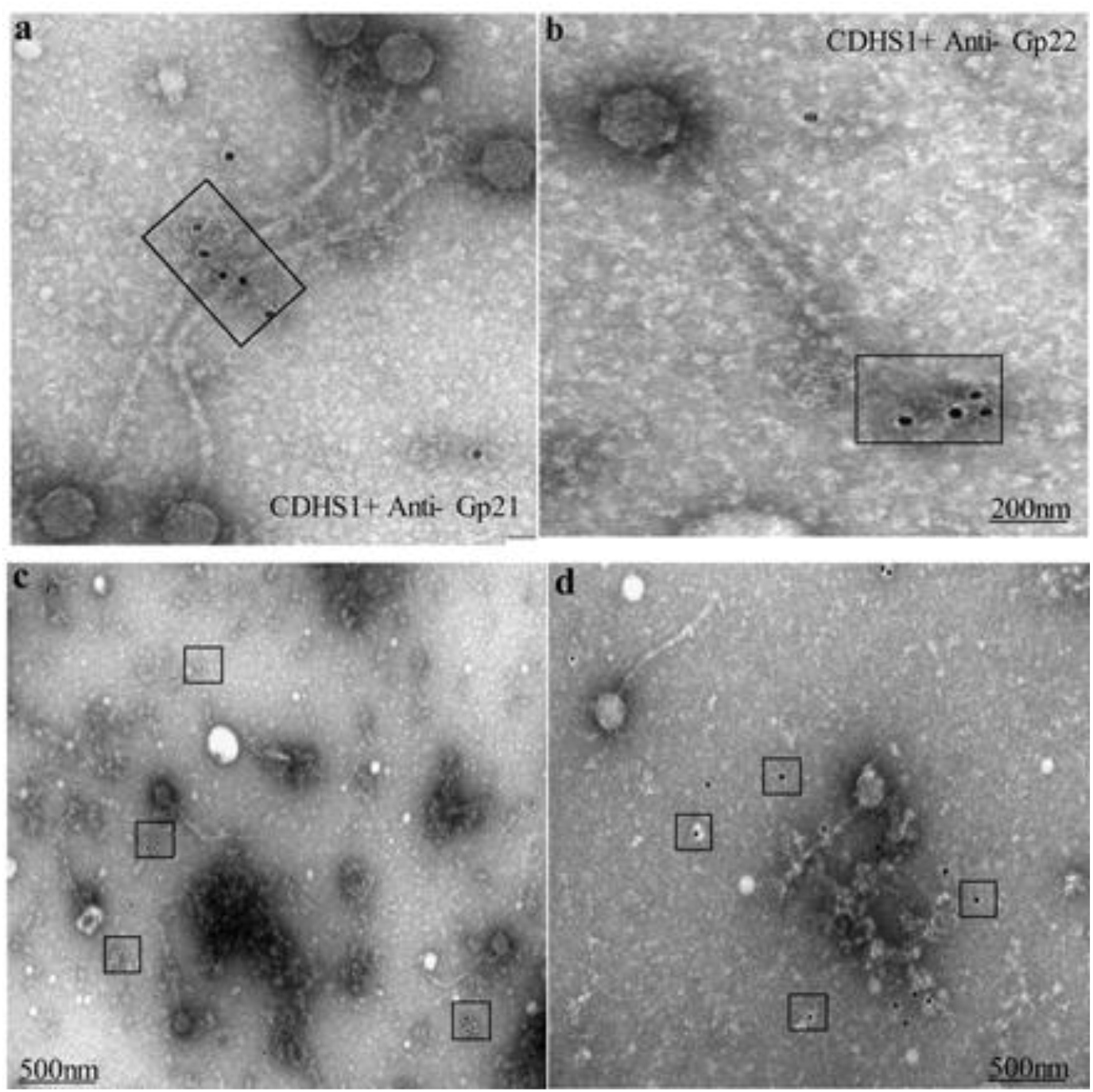
Immunogold labelling of tail proteins, Gp21 (ALY06964.1) and Gp22 (ALY06965.1). Transmission electron microscopy (TEM) images of negatively stained phage CDHS-1 after immunogold labelling with anti-Gp21 (ALY06964.1) serum (A) and anti-Gp22 (ALY06965.1) serum (B). (C) and (D) are the negative controls (pre-immune sera for each antibody raised).

### Structure of Gp22 (ALY06965.1) (RBP)

To characterise Gp22 (ALY06965.1) further, recombinant protein was purified by affinity and gel filtration chromatography. Gel filtration analysis showed that Gp22 (ALY06965.1) is a stable dimer in solution with an apparent molecular mass of ~130 kDa (expected mass of a dimer 136.4 kDa) (Figure 4a). Crystals of selenomethionine-enriched Gp22 (ALY06965.1) were grown at pH7 in the buffer containing Mg^2+^ and diffraction data were collected at Diamond Light Source. Phases were solved by selenomethionine single-wavelength anomalous dispersion, and data diffracted to 2.5Å resolution (Supplementary Table 1). Four copies of Gp22 (ALY06965.1) were observed in the asymmetric unit, all forming dimers through association with adjacent polypeptides about the crystallographic symmetry axes. Each polypeptide comprises an N-terminal L-shaped α-helical superhelix domain (529 residues) and a C-terminal β-sandwich domain. The superhelix domain comprises 42 α-helices (Figure S6). α-Helices of the long and short arms of the L pack in a parallel conformation with three α-helices per turn to form the right-handed superhelix. The shorter arm comprises 11 α-helices of between 4 and 17 residues, separated by loops of between 1 and 10 residues. The longer arm comprises 29 helices of 5 to 14 residues, separated by short loops of 1 to 3 residues. A hairpin-like structure formed from an antiparallel two-helix coiled-coil of α-helices (helices α24 and α25; FigureS6 and 4b) is located in the middle of the superhelix of the longer arm and projects at right angles to the plane of the superhelix.

**Figure 4.**
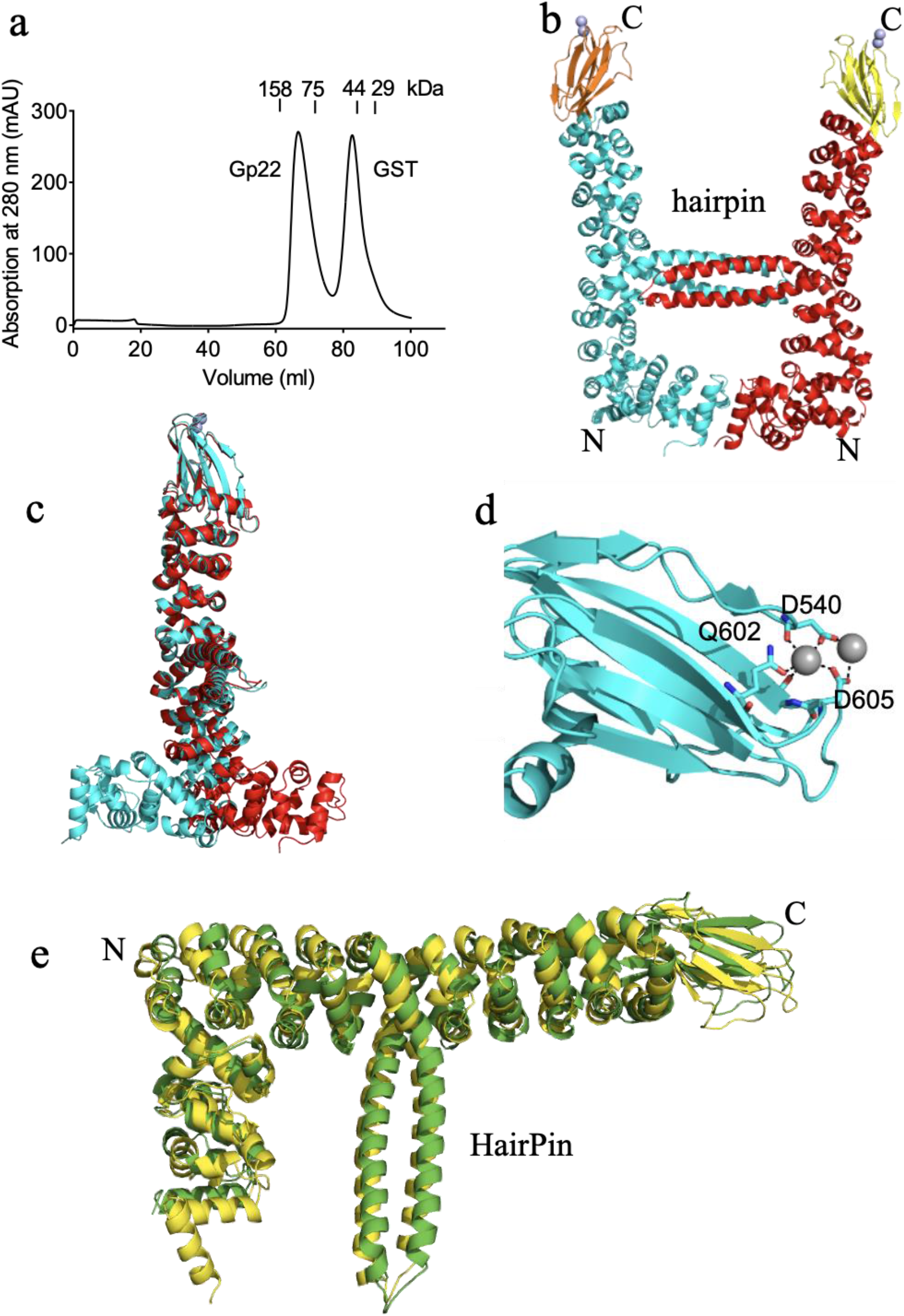
Gp22 (ALY06965.1) is a stable homodimer. a) gel filtration of the Gp22-GST fusion following cleavage by TEV protease. Gp22 (ALY06965.1) elutes with an apparent molecular mass of ~130 kDa from a Superdex 200 16/60 column based on the elution of molecular mass standards indicating that it is a dimer (monomer molecular mass = 68.4 kDa). GST is also dimeric as expected with a molecular mass of ~50 kDa b) the crystal structure of the Gp22 (ALY06965.1) dimer. The α-helical superhelix domains are in red, cyan, and the β-sandwich domains in yellow and orange. N- And C-termini are indicated c) overlay of the Gp22 polypeptides of the dimer. The β-sandwich domains and the C-terminal portions of the superhelix domains superpose closely, however the N-terminal portions are rotated by ~60° in opposite directions. d) the β-sandwich domain showing bound metal ions. Putative coordination ligands are indicated. e) Overlay of Gp22 (ALY06965.1) AlphaFold2 predictions (yellow) and Gp22 (ALY06965.1) crystal structure (green), AlphaFold2 accurately predicted Gp22 (ALY06965.1) structure, The main difference between the prediction and crystal structure is the angle between the N- and C-terminal arms, which is fixed by the dimer interface

The C-terminal domain comprises 8 antiparallel β-strands (β3 – β10; Figure S6) packed into two non-contiguous β-sheets. Strands β3, β10, β5 and β8 form one sheet and β7, β6, β9 and β4 the other. The domain contains two metal-binding sites in which the coordination geometry is compatible with Mg^2+^ present in the crystallization buffer. Coordination ligands include the side chains of Asp540, Gln602, and Asp605, and the main-chain carbonyl groups of Gly606 and Asp540 for one Mg^2+^ and the side chains of Asp540 and Asp605 for the second, more peripheral Mg^2+^ (Figure4 d). Caution is needed in assigning coordination ligands given that the data were only of medium resolution and only certain water molecules were observed in the density.

### Comparison between crystallographic resolution and AlphaFold 2 prediction

AlphaFold2 was used to predict the Gp22 (ALY06965.1) structure and the result was strikingly similar to the crystal structure. The RMSD for the short (residues Gln13-Thr158) and long (Ile159-Ile616) arms were 0.863 over 973 atoms and 1.50 over 2493 atoms, respectively (Figure 4e). The main difference between the prediction and crystal structure is the angle between the N- and C-terminal arms, which is fixed by the dimer interface. Notably, the junction between the β-sheet and superhelical domains and the alignment of the hairpin relative to the superhelical domain predicted correctly (Figure 4e). To test the accuracy of the structure of Gp22 (ALY06965.1) predicted by AlphaFold2, we used a C-terminal fragment (residues 358-616) as a search model for molecular replacement using the Phaser software package. The phases were solved using the default settings, with all four copies of the fragment correctly positioned in the asymmetric unit.

### The role of Mg^2+^ in CDHS-1 phage infection

Many phages require divalent cations ions to infect their bacterial hosts. To determine if divalent cations have a role in the CDHS-1 infection, phage was added to CD105LC1 in the presence of Mg^2+^ and/or Ca^2+^. The data show that CDHS-1 could only infect when Mg^2+^ was present in the media (Figure S4).

### The S-layer proteins of *C. difficile* contain the receptors for CDHS-1

To identify if S-layer Slp A protein acts as a receptor for phage CDHS-1, CDHS-1 was added to lawns of the S-layer SLp A deficient mutant FM2.5. CDHS-1 could not infect this strain (Figure S5b) at any titer, whereas it infected the wild-type CDR20291 strain efficiently (Figure S5a). Thus, S-layer Slp A protein is likely to act as the receptor for CDHS-1.

## Discussion

The recent explosion in novel and diverse phage discovery has not been matched by a mechanistic understanding of their biology. This mismatch stems from the time taken to obtain sequence data is significantly less than that needed to functionally characterise proteins. Structure/function data is key to improving our understanding of phage-bacterial interactions and exploitation ^2^. Developing phage products for therapy requires us to minimise the development of phage resistance in the bacterial host. Resistance mechanisms include the prevention of phage adsorption and DNA injection, restriction enzymes to degrade phage DNA, and CRISPR/Cas systems ^34^. Of particular relevance to this work is the desire to minimise resistance due to phage attachment. A useful strategy to overcome receptor-based resistance is to incorporate several phages with different RBPs within a cocktail, so if resistance develops to one RBP, other phages are still active^35^. Therefore, understanding, and characterising RBP’s is a crucial component of phage therapy applications. Here we demonstrate using antibody, neutralisation, and imaging studies that Gp22 (ALY06965.1) is the RBP for CDHS-1 and essential for *C. difficile* infection.

RBPs have been characterized for several phages that target Gram-negative bacteria, including *E. coli* phages T4^16^, lambda and T5^36,1^. Relatively little is known about RBPs from phages that infect Gram-positive bacteria and less still for *C. difficile* phages. For all known *L. lactics* siphoviruses RBPs, are trimeric complexes, with each monomer having a modular organization consisting of head, neck, and shoulder domains, in which the shoulder domain contains the binding site for the bacterial host^5,37,38^. *S. aureus* φ11 RBP again has three parts, a stem, platform, and tower domain^38^.

In contrast, Gp22 (ALY06965.1) is unlike any previously structurally characterised RBPs, representing a new RBP class. The Gp22 (ALY06965.1) is a U-shaped homodimer stabilised via a central crossbar formed from a four-helix bundle. Each monomer comprises an N-terminal L-shaped α-helical superhelix domain and a C-terminal β-sandwich domain. The protomers of each dimer are asymmetrical with the short arms rotated in opposite orientations to form the base of the U-shape. Gp22 (ALY06965.1) was crystallised in the presence of Mg^2+^ cations, compatible with our finding that Mg^2+^ is bound at the tip of the β-sandwich domain. This observation likely explains the observation CDHS-1 binding is Mg^2+^-dependent, although other mechanisms are possible.

We also used the protein structure prediction algorithm AlphaFold2 to predict the Gp22 (ALY06965.1) structure, which closely matched experimentally derived Gp22 (ALY06965.1) structure. The significance of applying such high-quality prediction software to phage proteins, specifically RBPs, will expand our ability to identify proteins with no sequence similarity to existing proteins but that have commonality in structure.

To determine what Gp22 (ALY06965.1) bound to, we showed that CDHS-1 could not infect the S-layer protein A mutant FM2.5, indicating that the CDHS-1 receptors are the S-layer protein SLpA. The FM2.5 strain has a point mutation at *SlpA* gene resulting in lack of the S-Layer protein (SlpA). However, minor cell wall proteins including Cwp2 and Cwp6 are still expressed^25^. S-layer protein SlpA was found as a receptor for Avidocin-CDs. The specificity of Avidocin-CD for *C. difficile* was acquired from fusing predicted RBPs from phages including phi-027b, phi-123, phi147, phi-242.c1 and Phi-68.4^25^. There is no sequence similarity between Avidocin-CD RBPs and CDHS-1 Gp22 (ALY06965.1).

The majority of phages infecting Gram-positive bacteria for which bacterial surface receptors have been identified, have receptors within two main components on the surface of Gram positive bacteria; teichoic acid or peptidoglycan^39^.

This is the first study to identify the RBP structure from any *C. difficile* phage. The results obtained establish a solid basis to understand how this phage attaches to *C. difficile*. This structural information will further the development of sensitive affinity-based infection diagnostics and therapeutics for this organism. Similar studies are ongoing for myoviruses that infect *C. difficile*, which we will report on in due course.

Importantly this work changes the paradigm of existing phage receptor binding proteins as belonging to known structural families: one of the key messages from the study is that there are likely to be completely novel classes of receptor binding proteins in existence which can only be identified through this combination of biochemical and structural biological approaches. Major limiting steps in functional characterisation are determining which genes encode for proteins of interest when there are no structural similarities with other phages and by technical difficulties of characterising phage proteins that are often particularly difficult to express, crystalise and diffract. The AlphaFold2 prediction of the CDHS-1 structureome confirmed the structure of proteins we expected to find within the tail unit and suggested a structure for the protein our phenotypic studies suggested was the RBP. These data suggest that using state-of-the-art machine learning for routine structural predictions can free us from dependence on sequence homology, which is sorely lacking in the known phages, and will significantly improve the speed by which we can understand the biology of phages. Our data shows that AlphaFold2 will be very useful to guide these studies and may indeed radically transform the speed by which novel families of phage receptor proteins can be identified and thus speed up practical applications such as phage therapy.

## Methods

### Bacteria, phages, and culturing

The *C. difficile* strain used in this study for propagating phage CDHS-1 is CD105LC1 of ribotype 027, in addition to CDR20291 of ribotype 027, They both were from our Laboratory collections, and they have been previously described in detail ^12,40^. Moreover, FM2.5 SlpA knockout (truncate SlpA at a site N-terminal to the post-translational cleavage site and, thereby, prevent the formation of an S-layer) derivative of CDR20291. *C. difficile* culturing and phage propagation were carried out according to a previously published protocol (Shan *et al*., 2012). Briefly, *C. difficile* was grown under anaerobic conditions (10% H_2_, 5% CO_2_, and 85% N_2_) on Brain-Heart Infusion (BHI, Oxoid, Basingstoke, UK) 1% agar plates, supplemented with 7% defibrinated horse blood (DHB, TCS Biosciences Ltd., Buckingham, UK) for 24 hours at 37°C. Liquid cultures were prepared by taking a single colony from the blood agar plate and then inoculated in a bijou tube containing 5 ml of Fastidious Anaerobic Broth (FAB; BioConnections, Kynpersley, UK). The liquid cultures were left to grow anaerobically overnight. Then, 500 μl of overnight FAB culture was inoculated into 50 ml of pre-reduced BHI and incubated until an OD550 of 0.2 was reached. Subsequently, 500 μl of phage CDHS-1was added to the culture and incubated for an additional 24 hours, followed by centrifuging at 3,400-x *g* for 10 min. The resulting supernatant was filtered using a 0.22 μm filter, and the phage titer in the filtrate was determined using a spot test assay according to a peer-reviewed protocol^12^.

### Protein Expression and Purification of the baseplate proteins Gp21(ALY06964.1) & Gp22 (ALY06965.1) for CDHS-1 phage

DNA extraction was carried out as previously described (Nale *et al*., 2016). Post DNA extraction, both *gp21* & *gp22* were amplified using PCR. The primers used for the PCR are 5-GTGATAAATTTGAGAGATAG-3 and 5-TTAACTCACCTCTTCTTTTATTTC-3 targeting *gp21gene*, and 5-TACTTCCAATCCATGAGTTGGGCGGAGACATACAAAG-3 and 5-TATCCACCTTTACTGTCATTAAATTGCTTGATACATTGCGTAA-3 to amplify *gp22* gene. Then, the amplified genes (*gp21* and *gp22*) were cloned into pET-based expression plasmids with the help of the cloning service (PROTEX) based at the University of Leicester. The resulting plasmids were used to transform *E. coli* BL21 (DE3) and an established protocol of protein expression in *E. coli* using isopropyl β-D-1-thiogalactopyranoside (IPTG) as inducer was followed (Campanacci *et al*., 2010). The proteins were purified using Affinity chromatography purification on Glutathione Sepharose 4B beads affinity column (GE Healthcare). After that, the proteins were further purified by gel filtration on 200 16/60 columns (GE Healthcare). In 20 mM Tris pH 7.5, 20 mM NaCl and, then concentrated by filtration using a 10-kDa molecular mass cut off the membrane (Amicon) before further usage.

### Phage Neutralisation

To determine which phage CDHS-1 tail protein was responsible for binding to *C. difficile*, the purified Gp21(ALY06964.1) and Gp22 (ALY06965.1) proteins were sent to Eurogentec (Brussels, Belgium) for the generation of polyclonal antibodies. The resulting anti-serum antibodies were used in a phage neutralization test to determine which of these anti-sera would be able to neutralize the phage infection. The assay was carried out according to a published protocol ^23^. Briefly, each antibody serum was diluted using SM buffer (10 mM NaCl, 8 mM, MgSO_4_.7H_2_O, and 50 mM Tris-HCL pH 7.5), into 1:10, 1:100, 1:1000 and 1:10000. Then 10^5^ phage CDHS-1 was added to each of the dilutions mentioned above. The mixture was incubated for 20 minutes at 37 °C. Then the mixture was serially diluted using SM buffer and spotted on the lawn of the *C. difficile* strain used.

### The interaction of Gp22 (ALY06965.1) protein with CD105LC1 strain

To determine if the recombinant Gp22 (ALY06965.1) interacts with CD105LC1, an adsorption inhibition assay was performed as described in^40^. Briefly, a strain grown in anaerobic conditions. Phage CDHS-1 (10^7^ PFU/ml) and different concentrations of Gp22 (ALY06965.1) proteins of 400μg, 200μg, 50μg, 0μg were added into CD105LC1 culture with an OD_600_ of around 0.2, respectively. After an incubation time of 30 minutes at 37°C, a spot test was carried out to determine the phage titer. To visualize the direct interaction between Gp22 and CD105LC1 Glutathione Sepharose 4B beads coated with either GST-tagged Gp22 (ALY06965.1) protein or only GST. Then they were mixed with CD105LC1 cultures and incubated at 4°C for 30 minutes before the confocal microscope. This assay was done as described in^41^.

### Immunogold labeling

Immunogold labeling was carried out to localize the phage tail protein Gp21(ALY06964.1) and Gp22 (ALY06965.1) on the phage particle. The assay was done as described previously with slight modification^23^. In an Eppendorf tube, each antibody serum was diluted 1:100 using SM buffer and then incubated with 10^9^ PFU/ml CDHS-1 phage for 20 minutes at room temperature. The mixture was then applied to glow discharged carbon-coated grids followed by grid washing with SM buffer for 10 minutes. Then a 1:30 diluted goat anti-rabbit IgG coupled with 12 nm gold colloids (Dianova, Hamburg) was added onto the grids and left for 20 minutes. Finally, the grids were negatively stained with 1% (w/v) uranyl acetate before examination under the TEM.

### AlphaFold2 Predictions

All 53 predicted protein sequences were used as input for the AlphaFold2 predictor as described (https://github.com/deepmind/alphafold) on the Danish National Supercomputer Computerome, using 20 NVidia Tesla VT100 GPUs. The pLDDT scores and highest ranked structures were extracted from the ranked_0.pdb and ranking.json result files and visualised with pymol^42^(The PyMOL Molecular Graphics System, Version 2.0 Schrödinger, LLC; https://pymol.org/2/)

### Gp22 (ALY06965.1) crystallization

To determine Gp22 (ALY06965.1) structure, selenomethionine labeling was performed, using inhibition of the methionine pathway to increase incorporation of the labelled methionine^43^.

Briefly, *E. coli* BL21 (DE3) transfected with the plasmid was grown overnight in 5 ml of LB. Then the cells were centrifuged for 5 minutes at 1300 x *g*. After that, the pellet was resuspended in 1 ml of M9 (Molecular Dimensions, UK) (M9 prepared according to the manufacturer’s guide) and added to 1 L of the same medium (M9). Cells were left to grow in a 37°C shaking incubator until they reached OD_600_ of 0.2, then amino acids were added; lysine, phenylalanine, and threonine at a final concentration of 100 mg/ml, and isoleucine, leucine, and valine at a final concentration of 50 mg/ml. Thereafter, the cells were grown until an optical density OD_600_ of 0.4 at 37°C, IPTG was added at a concentration of 0.5 mM to induce protein expression; finally, cells were incubated at 17°C overnight. Selenomethionine-Gp22 (ALY06965.1) protein purified as described above.

The purified Selenomethionine-Gp22 (ALY06965.1) protein was crystallized using the sitting-drop vapour diffusion technique. The crystals were grown in 0.1M HEPES pH 7, containing 0.05M ammonium acetate, 0.15M magnesium sulphate heptahydrate, 12% PEG smear medium, and 4% acetone. Then the crystals were transferred to a reservoir solution containing 30 % glycerol as a cryo-protectant before freezing in liquid nitrogen. The diffraction data were collected at the Diamond light source.

## Acknowledgments

Thanks, the ministry of higher education, Libya for my PhD sponsorship, Thanks to Marco Oggioni for for access to FM2.5 mutant. Thanks to Thanks to Natalie Allcock from University of Leicester for producing Transmission Electron Microscopy Images. Thanks to Mohammed Madadha for his technical support.

## Authors Contributions

Ahmed S. A. Dowah, Russell Wallis, Guoqing Xia and Martha R. J. Clokie designed the experiments. Ahmed S. A. Dowah performed the cloning, protein expression and purification experiments, Neutralization assay and Characterisation of the CDHS-1 tail module, Ahmed S. A. Dowah and Russell Wallis performed Gp22 Crystlisation and solved Gp22 crystal structure. Andy Millard, Bent Petersen, Thomas Sicheritz-Pontén and Martha R. J. Clokie performed Alpha Fold2 predictions of Gp22 and CDHS-1 structurom. Ahmed S. A. Dowah and Ali Abdul Kareem Ali performed the TEM Localisation of Gp22 and Gp21. Ahmed S. A. Dowah drafted the manuscript. Anisha M. Thanki, Jinyu Shan, Guoqing Xia, Andy Millard, Bent Petersen, Thomas Sicheritz-Pontén, Russell Wallis and Martha R. J. Clokie revised and edited the manuscript.

