## Supplementary Materials for "The structurome of a *Clostridium difficile* phage and the remarkable accurate prediction of its novel phage receptor-binding protein"

### Supplementary Data.

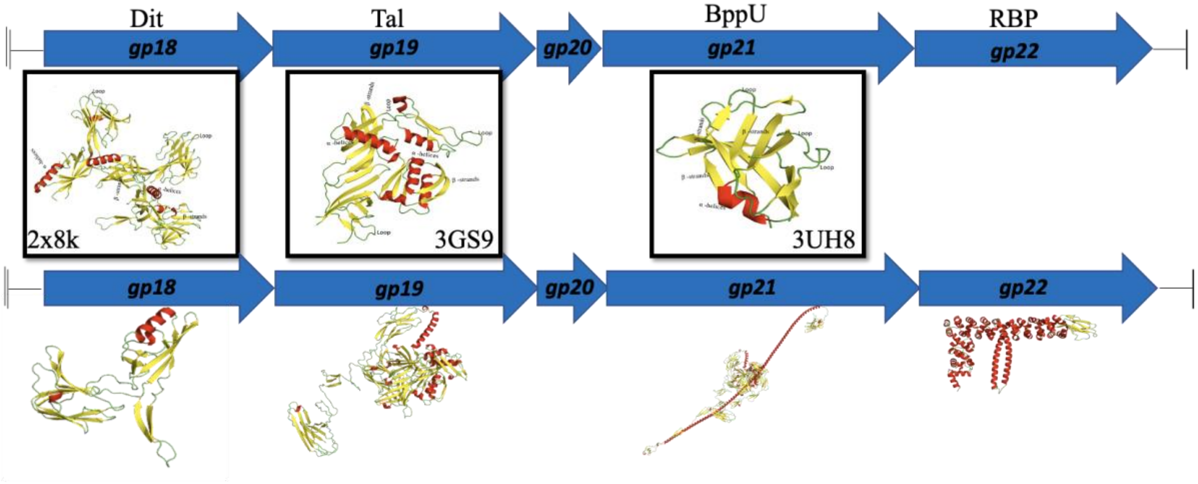


**Figure S1: HHpred analysis of the four structural gene products following the *tmp* gene of phage CDHS-1.**

Arrows represent genes *gp18*, *gp19*, *gp21*, *gp22*. The tail proteins encoded by these genes are indicated above the arrows. The structural homologues of these tail proteins are presented in the boxes beneath the corresponding genes. The PDB identifiers and ribbon structures (α-helices in blue, β-strand are shown for the structural homologues, and indicated by the boxes. The PDB entries shown here include 2X8K, *Bacillus* phage SPP1 baseplate Dit protein; 3GS9, tail associated lysin (Tal) of Gp18 (NP_465809.1) from *L. monocytogenes* phage A118 and 3UH8, is the N-terminal of BppU of *Lactococcus* phage TP901-1. AlphaFold2 predicted Gp18 (ALY06961.1) and Gp19 (ALY06962.1) structure accurately similar to HHpred analysis, However, the AlphaFold2 predicted the complete structure of Gp21(ALY06964.1) only the N-terminal was similar to HHpred prediction, and strikingly predicted Gp22 (ALY06965.1) which identical to Gp22 (ALY06965.1) crystal structure.

```
CDHS1_22       1 MSWAETYKVNSDLQGEPLNFLSYLQDIKLNGLDSYVLFIGNARIWEELYLNSLYLFSDRG
phiCD38.2      1 MSWAETYKVNSDLQGEPLNFLSYLQDIKLNGLDSYVLFIGNARIWEELYLNSLYLFSDRG
phiCD146       1 MSWAETYKVNSDLQGEPLNFLSYLQDIKLNGLDSYVLFIGNARIWEELYLNSLYLFSDRG
phiCD111       1 MSWAETYKVNSDLQGEPLNFLSYLQDIKLNGLDSYVLFIGNARIWEELYLNSLYLFSDRG

CDHS1_22      61 IRETVYTAFSETDIDNLFNKSTKLGEQLNAFYRTDIFSLGNADNVVKEMTIEHYNSLEEK
phiCD38.2     61 IRETVYTAFSETDIDNLFNKSTKLGEQLNAFYRTDIFSLGNADNVVKEMTIEHYNSLEEK
phiCD146      61 IRETVYTAFSETDIDNLFNKSTKLGEQLNAFYRTDIFSLGNADNVVKEMTIEHYNSLEEK
phiCD111      61 IRETVYTAFSETDIDNLFNKSTKLGEQLNAFYRTDIFSLGNADNVVKEMTIEHYNSLEEK

CDHS1_22     121 FKAGYDRYVTREQEKSTIGAWFNSTFSLDNTDLENLTTIEEILANVEATNAILNNSNAIV
phiCD38.2    121 FKAGYDRYVTREQEKSTIGAWFNSTFSLDNTDLENLTTIEEILANVEATNAILNNSNAIV
phiCD146     121 FKAGYDRYVTREQEKSTIGAWFNSTFSLDNTDLENLTTIEEILANVEATNAILNNSNAIV
phiCD111     121 FKAGYDRYVTREQEKSTIGAWFNSTFSLNNTGLESLTTIEEILANVEATNAILNNSNAIV

CDHS1_22     181 ALTMCKSSMDAVVASSNAMDLLGQYILRVTTESPVIRAILKNNVIRDAIINSDEAMTQIS
phiCD38.2    181 ALTMCKSSMDAVVASSNAMDLLGQYILRVTTESPVIRAILKNNVIRDAIINSDEAMTQIS
phiCD146     181 ALTMCKSSMDAVVASSNAMDLLGQYILRVTTESPVIRAILKNNVIRDAIINSDEAMTQIS
phiCD111     181 ALTMCKTSMDAVAASPNAMDLLGQYILKVTAEPAVIRAIIKNEVIRDAIINSDEAMTQIS

CDHS1_22     241 SNENSVMEIFNDLEATKVLVQNQNSINKILTNNVTVEKIIPNLLEMKYNLQTSLNYINTI
phiCD38.2    241 SNENSVMEIFNDLEATKVLVQNQNSINKILTNNVTVEKIIPNLLEMKYNLQTSLNYINTI
phiCD146     241 SNENSVMEIFNDLEATKVLVQNQNSINKILTNNVTVEKIIPNLLEMKYNLQTSLNYINTI
phiCD111     241 SNENSVMEIFNDLEATKVLVQNQNSINKILTNNVTVEKIIPNLLEMKYNLQTSLNYINTI

CDHS1_22     301 KSNIASGKGQIMAITYNEEIFPILKNAVKNYDGMETTRNISQRDIEEKIKISDAILESSI
phiCD38.2    301 KSNIASGKGQIMAITYNEEIFPILKNAVKNYDGMETTRNISQRDIEEKIKISDAILESSI
phiCD146     301 KSNIASGKGQIMAITYNEEIFPILKNAVKNYDGMETTRNISQRDIEEKIKISDAILESSI
phiCD111     301 KSNIASGKGQIMAITYNEEIFPILKNAAKNYDGMENTRNILQRDIEEKIKISDAILESSI

CDHS1_22     361 AMATFANNSIIVNKVGDRVGIIESIFSKTVSLNAFMKSTTAINILVNKTTAFTKIANNST
phiCD38.2    361 AMATFANNSIIVNKVGDRVGIIESIFSKTVSLNAFMKSTTAINILVNKTTAFTKIANNST
phiCD146     361 AMATFANNSIIVNKVGDRVGIIESIFSKTVSLNAFMKSTTAINILVNKTTAFTKIANNST
phiCD111     361 AMTTFANNSIIVNKVGDRAGIIESILGQTVPLNAFMKSTTAIKVLVNKATAFTKIVNNST

CDHS1_22     421 AFNAMLTISENNVTIANNTTAMGIIANNAQAMSTVANNDTSISVFVNNTTAMGIIA----
phiCD38.2    421 AFNAMLTISENNVTIANNTTAMGIIANNAQAMSTVANNDTSISVFVNNTTAMGIIA----
phiCD146     421 AFNAMLTISENNVTIANNTTAMGIIANNAQAMSTVANNDTSISVFVNNTTAMGIIA----
phiCD111     421 AFNAMLTISGNISSVANSSTAMNIIANNAQVMATVARNDTVLTVFINASTAITAIAGNVT

CDHS1_22     477 ------NSSTAMTKITLTGLALNRMVKSNTAKSILISKNSTLQTYKNNIQNTIQGSTAYF
phiCD38.2    477 ------NSSTAMTKITLTGLALNRMVKSNTAKSILISKNSTLQTYKNNIQNTIQGSTAYF
phiCD146     477 ------NSSTAMTKITLTGLALNRMVKSNTAKSILISKNSTLQTYKNNIQNTIQGSTAYF
phiCD111     481 SMSAIVGYSGAMNKITTSAFALNRIFKSTIGKDILISNNAILQTYKNNIHNTLRSSSGQF

CDHS1_22     531 RTITGFADADDNPPQTINSTYVGITYCYGYKGN-SYYGIVYHGYNTSIEAGRGNGYKDET
phiCD38.2    531 RTITGFADADNNPPQTINSTYVGITYCYGYKGN-SYYGIVYHGYNTSIEAGRGNGYKDET
phiCD146     531 RTITGFADADNNPPQTINSTYVGITYCYGYKGN-SYYGIVYHGYNTSIEAGRGNGYKDET
phiCD111     541 RLITSPTDSDSAPTQIINSSYVGVTYCYGYRGGTGKFAIVYHGHKTSAEAGRGDGGKDET
```

```
CDHS1_22     590  KKFITLGGARYDQSGDGYFTYAMYQAI
phiCD38.2    590  KKFITLGGARYDQSGDGYFTYAMYQAI
phiCD146     590  KKFITLGGARYDQSGDGYFTYAMYQAI
phiCD111     601  KKFITLGGTKFVESGDGYFNYSIYQAI
```

**Figure S2**: Alignment of Gp22 (ALY06965.1) protein sequences (RBps) of *C. difficile* phage CDHS-1, with other *C. difficile* phages PhiCD38.2, PhiCD146 and phiCD111. The alignment tools T-coffee: [T-Coffee Server] and Boxshade: [BoxShade Server] were used. The length of each sequence was 590 in the phages CDHS-1, PhiCD38.2 and PhiCD146. However, 601 for PhiCD111. The color of the amino acid represents its percentage identity to the consensus sequence, the black color indicates the higher the percentage identity. Gp22 (ALY06965.1) of CDHS-1 is conserved in phiCD38.6 and PhiCD146. However, it is varying towards the C. terminal of PhiCD111.

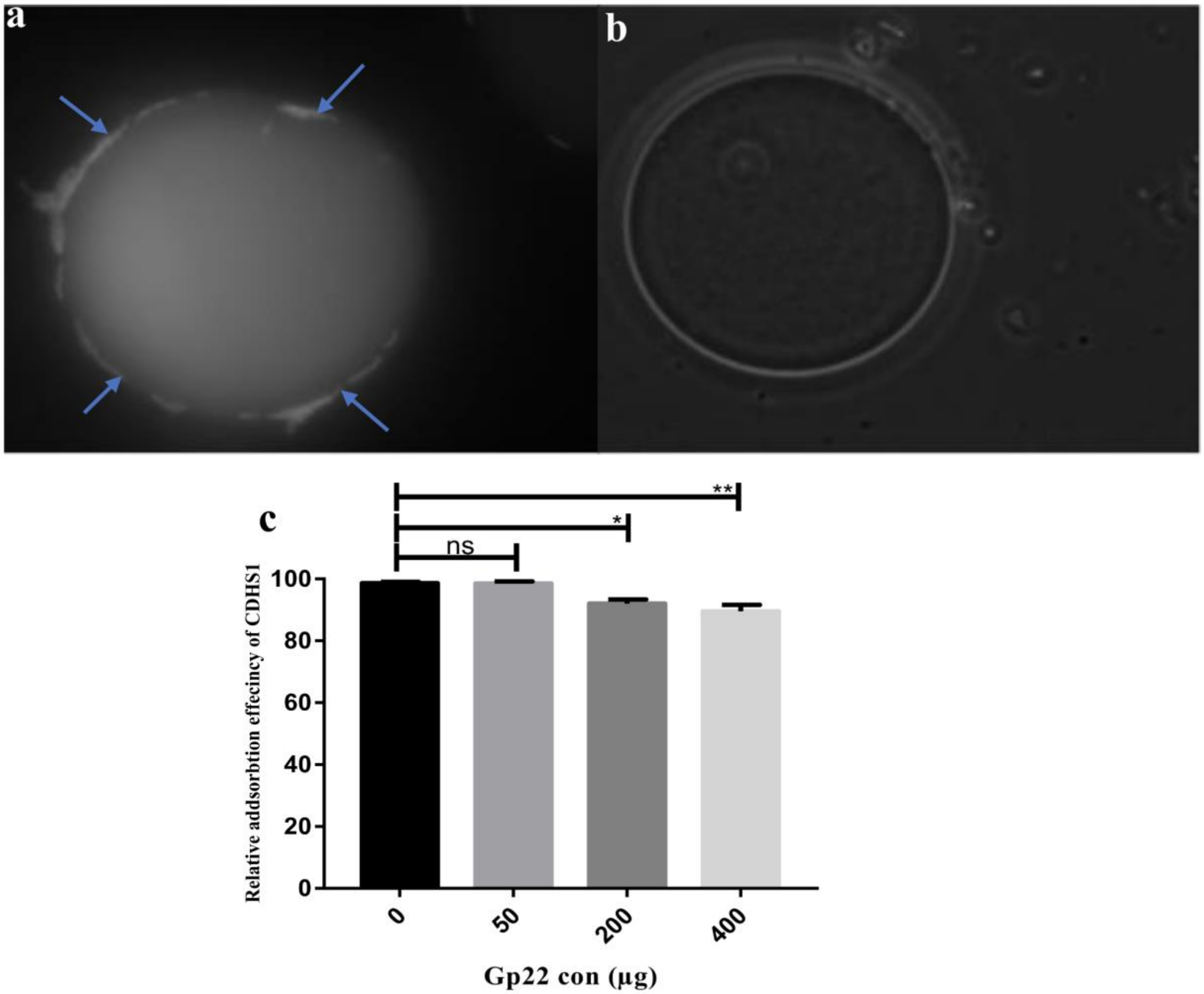


**Figure S3: The attachment of Gp22 (ALY06965.1) protein with CD105LC1 strain.**

(a) is a Confocal microscope image taken post-incubation of CD105LC1 strain with the Glutathione sepharose beads coated with Gp22 (ALY06965.1) protein tagged with GST. The result shows that the CD105LC1 cells attached to the beads coated with Gp22 (ALY06965.1) tagged with GST and indicates the Gp22 (ALY06965.1) protein has a role in phage binding with *C. difficile*. As a negative control, the CD105LC1 cells pre-incubated with Glutathione sepharose beads coated with GST only and the result shows no attachment in image (b). (c) is a Dose-dependent inhibition of CDHS-1 adsorption with recombinant Gp22 (ALY06965.1).

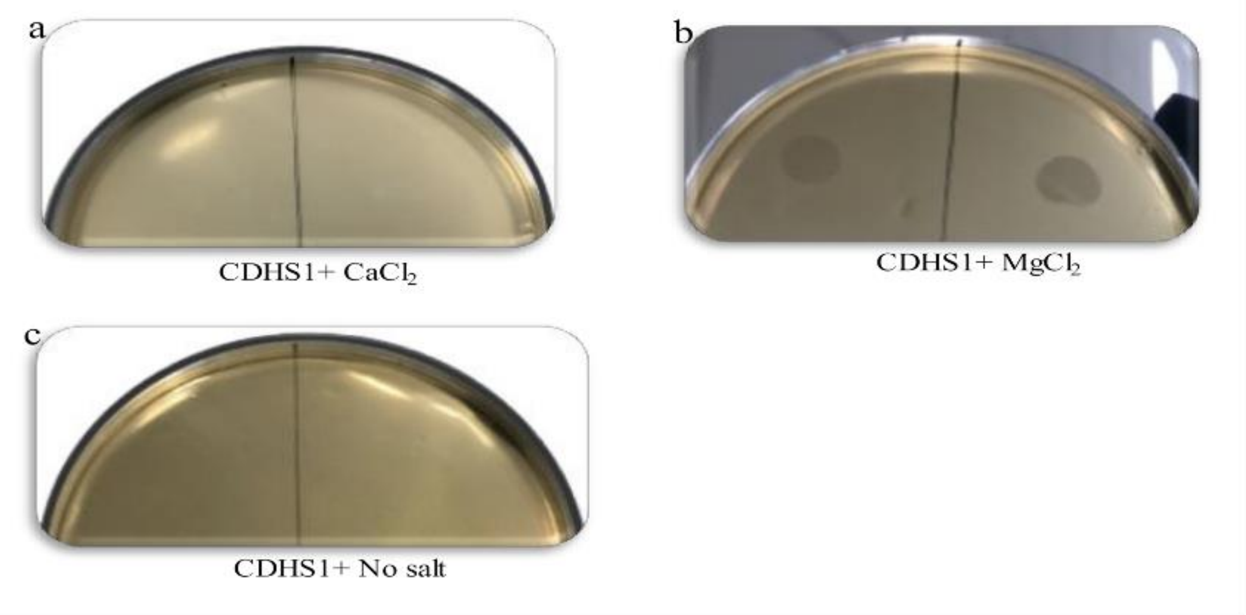


**Figure S4: the impact of Magnesium or Calcium on the CDHS-1 phage infection:** CDHS-1 spotted on a loan of CD105LC1 strain in the presence of $Ca^{++}$ (a) and $Mg^{++}$(b) and no salt (c) added to the media, the result indicates the CDHS-1 was able to infect only in the presence of $Mg^{++}$ (b). Indicating the CDHS-1 is a $Mg^{++}$ dependent for the infection.

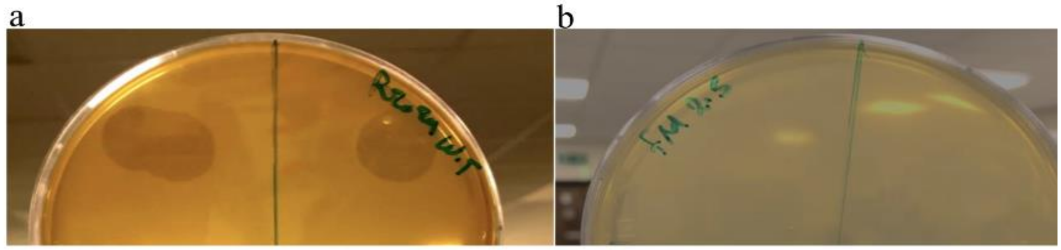


**Figure S5: CDHS-1 phage infection to CDR20291 and FM2.5 SlpA knockout derivative of R20291:** CDHS-1 spotted on a loan of CDR20291(a) and FM2.5 (b)strains, CDHS-1 was not able to infect the FM2.5 (b). As it did when spotted on the wild type CDR20291 (a). That indicates that receptors that CDHS-1 targeted are within the S-layer protein of CDR2029.

**Structure of Gp22 (ALY06965.1) (RBP)**

The total dimer interface covers 5523 $Å^2$ of surface, 2760 $Å^2$ from one polypeptide, and 2763 $Å^2$ from its partner. It is formed from interactions between the two α-helical hairpin-like structures that pack together to form a 4-helix bundle with 35 residues per helix. Additional contacts are made between the N-terminal portions of each superhelix domain. These domains come into contact through rotation of the short arms of the L-shaped polypeptides (by ~30°) in opposite orientations relative to the α-helical hairpin, thus creating an asymmetrical interface (Figure 4 c). Despite the asymmetry, the secondary structural elements of each protomer of the dimer are essentially the same, except that helix α16 is shorter in one of the protomers and disrupted at Tyr205 creating a kink (Figure S6) in the helix. The longer loop between helices α15 and α16 together with the kink in helix α16 allows the difference in the relative orientation of the shorter arms creating the asymmetry.

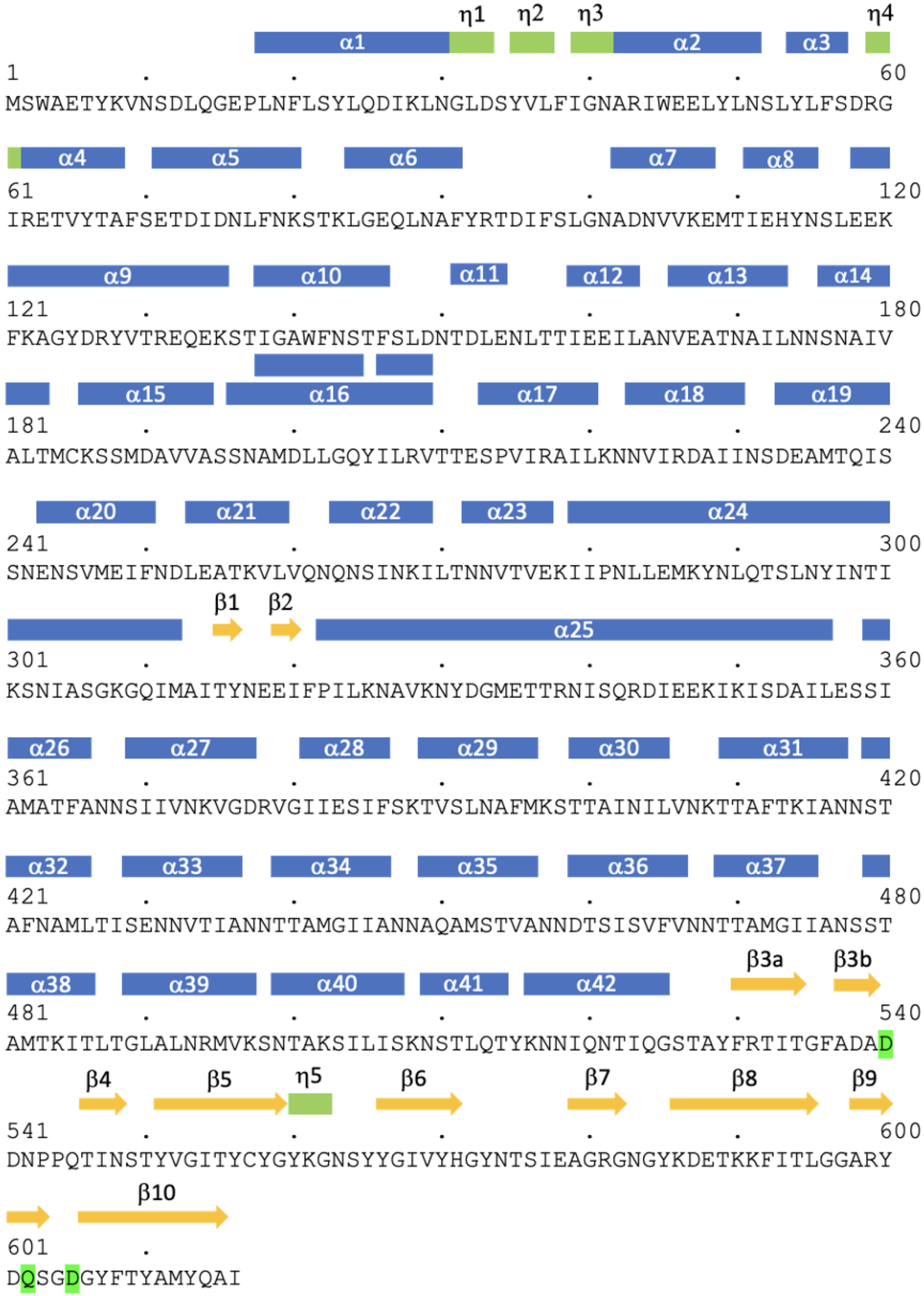
η1 η2 η3 η4
α1 α2 α3
1 . . . . . 60
MSWAETYKVNSDLQGEPLNFLSYLQDIKLNGLDSYVLFIGNARIWEELYLNSLYLFSDRG
α4 α5 α6 α7 α8
61 . . . . . 120
IRETVYTAFSETDIDNLFNKSTKLGEQLNAFYRTDIFSLGNADNVVKEMTIEHYNSLEEK
α9 α10 α11 α12 α13 α14
121 . . . . . 180
FKAGYDRYVTREQEKSTIGAWFNSTFSLDNTDLENLTTIEEILANVEATNAILNNSNAIV
α15 α16 α17 α18 α19
181 . . . . . 240
ALTMCKSSMDAVVASSNAMDLLGQYILRVTTESPVIRAILKNNVIRDAIINSDEAMTQIS
α20 α21 α22 α23 α24
241 . . . . . 300
SNENSVMEIFNDLEATKVLVQNQNSINKILTNNVTVEKIIPNLLEMKYNLQTSLNYINTI
β1 β2
α25
301 . . . . . 360
KSNIASGKGQIMAITYNEEIFPILKNAVKNYDGMETTRNISQRDIEEKIKISDAILESSI
α26 α27 α28 α29 α30 α31
361 . . . . . 420
AMATFANNSIIVNKVGDRVGIIESIFSKTVSLNAFMKSTTAINILVNKTTAFTKIANNST
α32 α33 α34 α35 α36 α37
421 . . . . . 480
AFNAMLTISENNVTIANNTTAMGIIANNAQAMSTVANNDTSISVFVNNTTAMGIIANSST
β3a β3b
α38 α39 α40 α41 α42
481 . . . . . 540
AMTKITLTGLALNRMVKSNTAKSILISKNSTLQTYKNNIQNTIQGSTAYFRTITGFADAD
β4 β5 η5 β6 β7 β8 β9
541 . . . . . 600
DNPPQTINSTYVGITYCYGYKGNSYYGIVYHGYNTSIEAGRGNGYKDETKKFITLGGARY
β10
601 .
DQSGDGYFTYAMYQAI

**Figure S6:** Sequence of Gp22 (ALY06965.1) showing the secondary structure elements. α-Helices are *in blue*, 310-helices (h) in *green* and β-strands in *yellow*. Differences in helix α16 in the two protomers of the asymmetric homodimer are indicated. Residues serving as putative coordination ligands for the two metal ions are shaded in *green.*

### Supplementary Tables

**Supplementary Table 1. Data collection and refinement statistics.**

| | |
|---|---|
| Data collection | |
| Beamline | Diamond Light Source I03 |
| Wavelength, *Å* | 0.9793 |
| Space group | P $2_1$ $2_1$ 2 |
| a, b, c, Å | 178.2, 227.5, 114.9 |
| α, β, γ, ° | 90, 90, 90 |
| Resolution, *Å* | 140.3 - 2.52 (2.82 - 2.52) |
| No. reflections | 104702 (5235) |
| R*sym* | 0.167 (1.07) |
| CC(1/2) | 1.0 (0.7) |
| I/δI | 9.1 (1.6) |
| Completeness | 95.8 (73.3) |
| Redundancy | 6.6 (7.1) |
| Resolution, *Å* | 140.3 - 2.52 (2.61 - 2.52) |
| No. reflections | 104684 (291) |
| Multiplicity | 6.6 (7.1) |
| R*work*/R*free* | 18.8/22.8 |
| No. atoms | 18877 |
| Protein | 18738 |
| Ligands | 8 |
| Water | 133 |
| B-factors, $Å^2$ | 46.4 |
| Protein | 45.7 |
| Ligand | 65.8 |
| Water | 40.5 |
| RMS (bonds) Å | 0.012 |
| RMS (angles) ° | 1.45 |

Statistics for the highest-resolution shell are shown in parentheses.

**Supplementary Table 2: confidence scores of AlphaFold2 prediction of the CDHS-1 structurome.**

| Accuracy | Mean pLDDT score | No of proteins | % | Annotation |
|---|---|---|---|---|
| High accuracy | >90 | 21 | 39.6% | hypothetical proteins, capsid proteins, tail protein, ParA, integrase, terminase, amidase, Cro, ssDNA binding proteins |
| Good accuracy | between 70 and 90 | 27 | 50.9% | |
| Low confidence | between 50 and 70 | 4 | 7.5% | conserved hypothetical proteins, tape measure protein |
| Unreliable | <50 | 1 | 1.9% | conserved hypothetical |
